# Low frequency of pathogenic allelic variants in the 46,XY differences of sex development (DSD)-related genes in small for gestational age children with hypospadias

**DOI:** 10.1101/748277

**Authors:** Barbara L. Braga, Nathalia L. Gomes, Mirian Yumie Nishi, Bruna Lucheze Freire, Rafael Loch Batista, Antonio Marcondes Lerario, Mariana Ferreira de Assis Funari, Elaine Maria Frade Costa, Sorahia Domenice, José Antonio D. Faria Junior, Alexander Augusto Lima Jorge, Berenice Bilharinho Mendonca

## Abstract

**Background:** Hypospadias is a congenital disorder of male genital formation where the urinary opening is not on the head of the penis and genetic factors play an important role in the incidence of this early developmental defect in 46,XY individuals, in both isolated and syndromic forms. Children born small for gestational age (SGA) present a high frequency of hypospadias of undetermined etiology, ranging from 15 to 30%, but the detection of hypospadias’ etiology remains low.

**Patients and methods:** from a cohort of 46,XY DSD patients, we identified 25 SGA children with medium or proximal hypospadias; four of them with associated syndromic characteristics. DNA samples from subjects were studied by massively parallel targeted sequencing (MPTS) using a targeted panel. MLPA was used for molecular diagnosis in two children with clinical phenotype of Silver Russel syndrome.

**Results:** Loss of DNA methylation (11p15 LOM) at ICR1 was identified in two out of four syndromic children. The other syndromic patient had 3M syndrome phenotype and two novel likely-pathogenic variants in compound heterozygous state were found in *CUL7* gene. The last syndromic subject had Mulibrey nanism and, a homozygous variant in *TRIM37* was identified in the patient and confirmed in heterozygous state in the mother. Among non-syndromic children seven rare heterozygous variants with uncertain significance in six DSD-related genes were identified: two children had *DHX37* variants associated with *GATA4* and *WWOX* variants, respectively; three children had heterozygous variants, in *WT1, IGF1R*, and *BMP8B* genes.

**Conclusion:** Pathogenic or likely-pathogenic variants in DSD-related genes were not identified in non-syndromic SGA children with hypospadias, suggesting that multi-factorial causes, unknown genes or unidentified environmental factors (epigenetic defects), may be involved in the etiology of this condition.

## Introduction

The child born small for gestational age (SGA) is defined as the one with birth weight and/or length 2 or more standard deviation (SD) below the population mean for gestational age. Approximately 10-15% of them do not recover growth in postnatal life. The underlying causes of pre and postnatal growth retardation are diverse including known genetic syndromes, such as Silver-Russell syndrome (SRS). However, hypospadias are presented even in cases that cannot be categorized in a particular syndrome (1,2). SRS is a clinical and (epi)genetic heterogeneous syndrome characterized primarily by pre and postnatal growth retardation. The most common molecular mechanisms are loss of methylation on chromosome 11p15 (11p15 LOM), occurring in 30-60% of patients. Some genotype-phenotype studies showed 11p15 LOM patients presenting more typical and severe presentations (3,4,5). Among the variable signs in patients with SRS, male genital abnormalities are presented in about 40% of the male patients with this syndrome (6). It is interesting to note that SGA patients also present a high frequency of genital atypia of undetermined etiology, ranging from 15 to 30% (7). Genetic factors play an important role in the incidence of hypospadias, but the detection of hypospadias’ etiology remains low (8). Our aim is to investigate the genetic cause underlining hypospadias in syndromic and non-syndromic SGA children.

## Materials and methods

Within a cohort of 272 subjects with 46,XY differences of sex development (DSD) followed in Clinical Hospital of University of São Paulo 59 have 46,XY DSD due to unknown cause. Among them, 30 are SGA children with medium or proximal hypospadias and were invited to participate in the study and 25 agreed to participate. Four of them had associated syndromic characteristics. This study was approved by the local medical ethics committee and patients and/or guardians gave their informed written consent. All patients have 46,XY karyotype in at least 30 analyzed cells obtained from peripheral blood. DNA samples from subjects and their families were obtained from peripheral blood leukocytes and were studied by massively parallel targeted sequencing (MPTS) using a targeted panel.

Two versions of the custom panel of target genes were designed using the *Agilent SureDesign 2*.*0 tool* (Agilent Technologies, Santa Clara, CA, USA) (Table 1). Genes that participate in the process of sex development and growth were included. Exonic regions of the target genes were included, including an area up to 25 bp before and after each exon.

**Table 1.** Genes included in both custom panels.

| Target genes |  |  |  |  |  |  |  |  |  |  |  |
| --- | --- | --- | --- | --- | --- | --- | --- | --- | --- | --- | --- |
| ACBD7 | ARX | BMP15 | CYP11A1 | DND1 | FST | HNF1B | KATNBL1 | NANOS3 | POR | SOX10 | SUPT3H |
| ACVR1B | ATRX | **BMP8B\*** | CYP17A1 | ESR1 | GADD45G | HPGDS | LEF1 | NCOA1 | PRKACG | SOX13 | TCF21 |
| ACVRL1 | BBS1 | BMPR1B | CYP19A1 | FAM189A2 | GATA4 | HSD11B1 | LGR5 | NCOA2 | PTGDS | SOX7 | TRIM32 |
| AKR1C2 | BBS10 | CAMK1D | CYP26B1 | FBLN2 | GATA6 | HSD17B3 | LHCGR | NCOA3 | PUM2 | SOX9 | **TRIM37** |
| AKR1C3 | BBS12 | CBLN1 | DHCR7 | FGD1 | GDF9 | HSD3B2 | LHX1 | NR0B1 | RSPO1 | SRA1 | UBE3A |
| AKR1C4 | BBS2 | CHD7 | DHH | FGF9 | GDNF | **IGF1R** | MAMLD1 | NR3C1 | RSPO2 | SRD5A2 | USP34 |
| AMH | BBS4 | CITED2 | DHX37 | FGFR2 | GPR56 | IHH | MAP3K1 | NR5A1 | SIX1 | SRY | WT1 |
| AMHR2 | BBS5 | CITED4 | DMRT1 | FOXL2 | GPR83 | INHBB | MKKS | PAPPA | SIX4 | STAG3 | WWOX |
| AR | BBS7 | CTNNB1 | DMRT2 | FRAT1 | GTF2F1 | IRX3 | MSX1 | PDGFA | SMAD4 | STAR | ZFPM2 |
| ARL6 | BBS9 | **CUL7** | DNAJC15 | FSHR | HHAT | KATNAL1 | NANOS2 | PDGFRB | SMARCE1 | STIM1 |  |

\*In bold, genes included only in the second version of the panel with significant variants.

The panel-based sequencing was performed in the Illumina MiSEQ platform (Illumina, Inc., San Diego, CA, USA). Paired-end reads were aligned to the hg19 assembly of the human genome with BWA-MEM. Variants were called and annotated with Platypus and ANNOVAR. Sanger sequencing was used to segregate the variants in the family. Multiplex Ligation-dependent Probe Amplification specific for Silver-Russel and Beckwith-Wideman syndromes (SALSA MLPA ME030 BWS/RSS) (MRC-Holland, Amsterdam, Netherlands) was used for molecular diagnosis in both children with SRS phenotype. The identified variants were classified according to American College of Medical Genetics (ACMG) criteria (9).

The targeted panel sequencing data were screened for rare variants (minor allele frequency < 0.1% in the public databases: Genome Aggregation Database (gnomAD) (10,11), 1000 Genomes (12), and in the Brazilian population database (ABraOM) (13), located in exonic and consensus splice site regions. Subsequently, the filtration pipeline prioritized potentially pathogenic candidate variants (loss of function variants and variants classified as pathogenic by at least three *in silico* programs).

The sequencing reads carrying candidate variants were visually confirmed using the Integrative Genomics Viewer (Broad Institute, Cambridge, MA). Candidate variants were segregated in the available family members by Sanger method.

## Results

### Syndromic SGA children

The 11p15 LOM was identified in two clinical SRS children by MLPA, confirming the clinical diagnosis of SRS. The other syndromic patient has 3M syndrome phenotype, and presents two likely pathogenic variants (p.Trp1622* and p.Gln897Hisfs*23) in compound heterozygosis state in *CUL7* gene. Sanger sequencing confirmed the heterozygous p.Gln813fs variant in his mother. The child is a SGA boy, born from non-consanguineous parents, with disproportionate short stature, proximal hypospadias, unusual facial features (hypoplastic midface; short, broad neck). The last syndromic subject is a SGA child, born from consanguineous parents, with triangular face, hypoplastic midface, frontal bossing, short stature, proximal hypospadias, feeding difficulties, and early respiratory distress. The homozygous p.Tyr341Ilefs*16 variant in *TRIM37* was identified in the patient. Sanger sequencing confirmed the heterozygous p.Tyr341Ilefs*16 variant in his mother (Table 2).

**Table 2.** Variants found in syndromic children with hypospadias

| Syndromic subject | Phenotype | Diagnosis | Gene | Variant | Protein |
| --- | --- | --- | --- | --- | --- |
| 1 | Growth failure, feeding difficulties, body asymmetry, protuding forehead, hypospadias | Silver-Russell syndrome | *H19* DMR | 11p15 LOM | - |
| 2 | Growth failure, feeding difficulties, body asymmetry, protuding forehead, hypospadias | Silver-Russell syndrome | *H19* DMR | 11p15 LOM | - |
| 3 | Proximal hypospadias, ocular hypertelorism, short nasal base, short neck, low cartilage ear pavement, small hands, single palmar fold, clinodactyly; difficulty growing and weight gain from birth | 3M syndrome | *CUL7* | c.4866G>A and c.2691delA | p.Trp1622* and p.Gln897Hisfs*23 |
| 4 | Proximal hypospadias, triangular face, frontal bossa, middle face hypoplasia, enlarged filter, articular foruxitude, hypospadias, and short stature | Mulibrey nanism | *TRIM37* | c.1020delC | p.Tyr341Ilefs*16 |

### Non-syndromic SGA children

Seven rare heterozygous variants with uncertain significance in six DSD-related genes were identified in five patients: *DHX37* variants (p. Val717Ile and p.Ala737Thr) were found associated with *GATA4* p.Pro407Arg and with *WWOX* p.Ttyr85Asp, respectively, in two children; three children each have heterozygous variants, the *WT1* p.Cys350Arg, the *IGF1R* p.Arg1337Cys, and the *BMP8B* p.Arg116Cys variant (Table 3). Sanger sequencing confirmed the occurrence of the heterozygous variant *BMP8B* p.Arg116Cys in the patient’s mother.

**Table 3.**
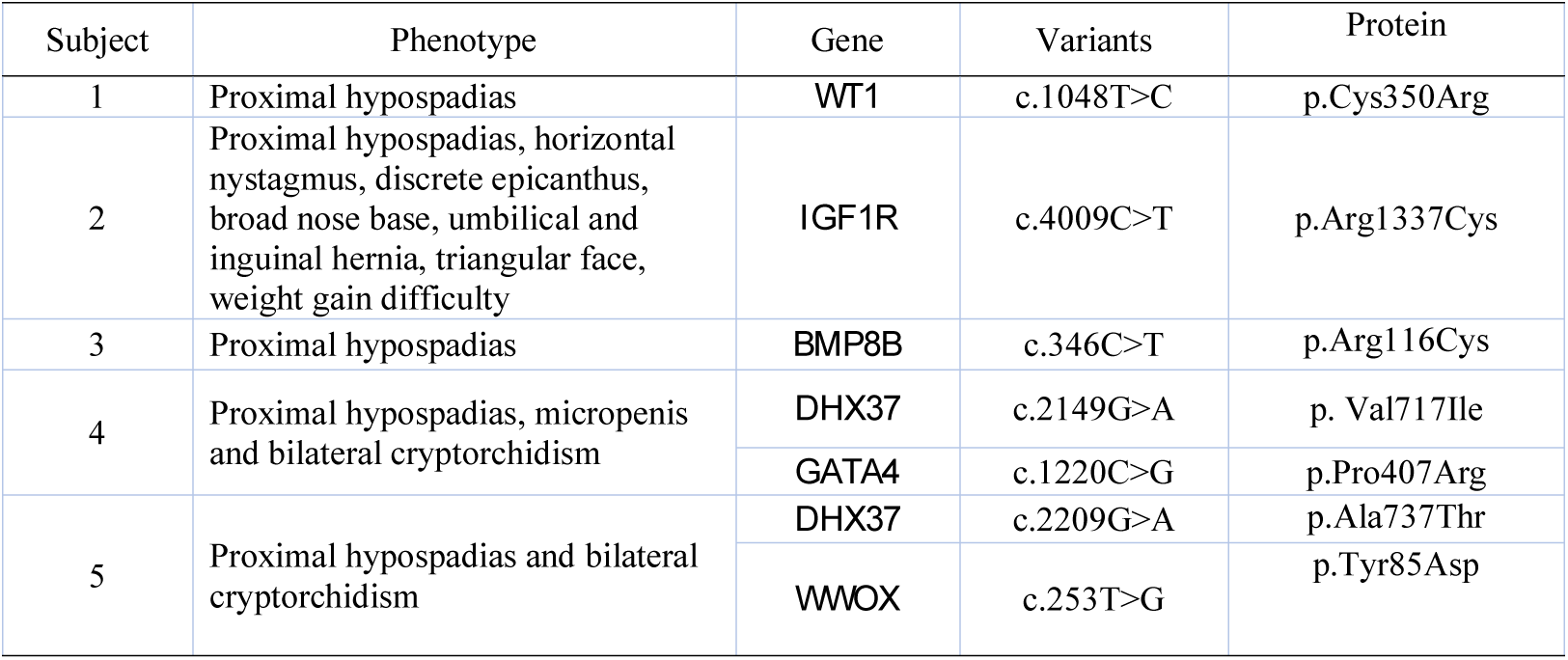
Uncertain significance rare heterozygous variants in SGA children with proximal hypospadias

## Discussion

SGA patients present a high frequency of hypospadias without a known etiology (8). Some studies believe that this low diagnosis could be explained because this condition has yet unknown genes and/or environmental factors involved (14, 15, 16). The present study investigated a possible molecular cause of hypospadias in syndromic and non-syndromic children.

In the syndromic SGA patients with hypospadias, the molecular diagnosis confirmed the clinical phenotype. Defects in *CUL7* gene are a known cause of 3M syndrome type 1, confirming the clinical features of the patient. The protein encoded by *CUL7* interacts with *TP53, CUL9*, and *FBXW8* proteins. *TRIM37* encodes a protein of the tripartite motif (TRIM) family, whose members are involved in diverse cellular functions such as developmental patterning and oncogenesis; mutations in *TRIM37* are associated with Mulibrey nanism, a serious autosomal recessive disorder. No defect in other gene associated with 46,XY DSD diagnosis was identified in the syndromic children.

SRS patients are characterized by a large spectrum of features and signs, varying from complex to milder phenotypes. Several clinical scoring systems have been proposed in the last few years. The First International Consensus Statement on diagnosis and management of SRS indicated the use of the Netchine-Harbison clinical scoring system (NH-CSS) for SRS clinical diagnosis. The most common molecular mechanisms are loss of methylation on chromosome 11p15 (11p15 LOM), occurring in 30-60% of patients, and maternal uniparental disomy for chromosome 7 (upd(7)mat), occurring in 5-10% of patients (3, 4, 5). Among the variable signs in patients with SRS, male genital abnormalities are presented in about 40% of the male patients with this syndrome (6). It is interesting to note that SGA patients also present a high frequency of genital atypia, ranging from 15 to 30%, of undetermined etiology (7). We hypothesized that genital atypia in SRS is not directly related to the epigenetic cause of the syndrome but it could share the same genetic mechanism of hypospadias in SGA children.

Studies of methylation role in specific disorders, as well as its influence on phenotypes would be helpful to increase our understanding about external genitalia development, contributing to expand the molecular etiology of hypospadias.

In non-syndromic SGA children, seven rare heterozygous variants with uncertain significance in six DSD-related genes were identified in these patients. The causes of hypospadias could be attributed to defects in genetic factors or considered syndrome-associated hypospadias (14, 17, 18). However, the association of these variants of uncertain significance with the phenotype could not be established in the absence of functional studies.

## Conclusion

In conclusion no proven genetic defects clarify the etiology of hypospadias in syndromic and non-syndromic SGA children, suggesting that multi-factorial causes, unknown genes or unidentified epigenetic defects, may be involved in the etiology of this condition.

